# X-linked palindromic gene families *4930567H17Rik* and *Mageb5* are dispensable for male mouse fertility

**DOI:** 10.1101/2021.11.10.468145

**Authors:** Evan R. Stark-Dykema, Eden A. Dulka, Emma R. Gerlinger, Jacob L. Mueller

## Abstract

Mammalian sex chromosomes are enriched for large, nearly-identical, palindromic sequences harboring genes expressed predominately in testicular germ cells. Discerning if individual palindrome-associated gene families are essential for male reproduction is difficult due to challenges in disrupting all copies within a gene family. Here we generate precise, independent, deletions to assess the reproductive roles of two X-linked palindromic gene families with spermatid-predominant expression, *4930567H17Rik* or *Mageb5*. Via sequence comparisons, we find mouse *4930567H17Rik* and *Mageb5* have human orthologs, *4930567H17Rik* is rapidly diverging in rodents and primates, and *4930567H17Rik* is harbored in a palindrome in humans and mice, while *Mageb5* is not. Mice lacking either *4930567H17Rik* or *Mageb5* gene families do not have detectable defects in male fertility, fecundity, spermatogenesis, or in gene regulation, but do show differences in sperm head morphology, suggesting a potential role in sperm function. We conclude that while all palindrome-associated gene families are not essential for male fertility, large palindromes influence the evolution of their associated gene families.

**Summary sentence:** Mice lacking X-palindromic gene families display normal male fertility, fecundity, spermatogenesis, and gene expression but exhibit differences in sperm head morphology, suggesting a potential role for these gene families in sperm development.

## Introduction

Large (>8 kb) palindromes are inverted segmental duplications that contain nearly-identical (>99%) DNA sequences. Large palindromes are enriched on the X and Y chromosomes in mammals [1, 2] and harbor gene families (≥ 2 copies of nearly-identical gene sequence) expressed predominantly during spermatogenesis [3, 4]. Males are more susceptible to deleterious mutations in single-copy sex-linked genes, because males are hemizygous (one X and one Y) [5]. Having a second copy of a sex-linked gene could potentially provide protection against hemizygotic susceptibility to new deleterious mutations. The presence of genes in two copies on either the X or Y in large palindromes may have evolved to provide protective roles for genes that are important for male fertility [6]. For example, in mice, three of four independent deletions of large palindrome arrays result in male infertility [7-9]. Additionally, large deletions of Y chromosome palindromes in men can result in male infertility [10, 11]. These previous findings suggest genes harbored within large palindromes may be essential for male fertility.

Deletions of large palindrome arrays also remove both associated gene families and the palindrome structures, making it difficult to separate whether the loss of the gene families or palindrome structures contribute to male infertility. Unlike palindrome arrays, which contain multiple segmental duplications in each arm, singleton palindromes contain a single segmental duplication in each arm. To investigate the importance of palindrome structure, previous studies have disrupted arms of singleton palindromes. Deletion or inversion of arms within singleton palindromes did not alter overall fertility [12]. These studies did not, however, investigate whether gene families within these palindromes are necessary for male fertility since one copy of the gene family was left intact within the remaining palindrome arm. If large palindromes provide protective functions for genes that are essential for male fertility, then complete loss of singleton palindrome-associated gene families could result in defects in spermatogenesis and male fertility.

To test if singleton palindrome-associated gene families are important for male fertility, we deleted both gene copies of the mouse X-linked gene families *4930567H17Rik* or *Mageb5*. These two gene families have predominant expression in spermatids and each gene family is the only known gene harbored within the arms of each singleton palindrome [12]. We find that both *4930567H17Rik* or *Mageb5* gene families in mice have orthologs in humans, but despite this conservation, mice lacking either gene family do not exhibit detectable defects in male fertility and post-meiotic spermatogenesis. We do observe several abnormalities in sperm head morphology, indicating that while *4930567H17Rik* or *Mageb5* are not necessary for male fertility, both genes likely play a role in post-meiotic sperm development. Overall, our study supports that gene families in singleton palindromes may play important roles in spermatogenesis, but are not always necessary for overall fertility. Our findings that *4930567H17Rik* or *Mageb5* are dispensable for male fertility is consistent with previous efforts demonstrating many single-copy testes-specific genes are also dispensable for male fertility in mice [13-15]. Our studies add to previous findings suggesting *4930567H17Rik* and *Mageb5* palindrome structures are not essential for male fertility or spermatogenesis [12].

## Materials and Methods

### Generation of mice lacking *4930567H17Rik* and *Mageb5* palindrome-associated gene families

Mice lacking the X-linked palindrome-associated *4930567H17Rik* and *Mageb5* gene families were generated using a CRISPR Cas9 strategy. We selected single guide RNAs (sgRNAs) within the coding sequences of *4930567H17Rik* or *Mageb5* (Table S1). The Cas9 (ESPCAS9PRO, Sigma-Aldrich/Merck KGaA, Darmstadt, Germany) cleavage efficiency of individual sgRNAs was determined via injection of sgRNA (30ng/ul) / Cas9 (50ng/ul) complexes into mouse zygotes and screening for edits via PCR using primers flanking sgRNA cut sites (Table S2) and subsequent Sanger sequencing. We selected sgRNAs with cleavage efficiencies of > 30% to delete *4930567H17Rik* and *Mageb5* gene families.

To generate mice lacking the *4930567H17Rik* gene family *(4930567H17Rik*^ΔCDS/Y^), C57BL/6J X SJL hybrid females were crossed with existing *4930567H17Rik*^+/Y^ mice [12]. Zygotes were injected with Cas9 protein (50 ng/μl), a single-stranded oligonucleotide donor (10 ng/μl), and dual sgRNAs (30 ng/μl) targeting each *4930567H17Rik* gene copy to achieve a ∼650 base pair deletion within each copy on both palindrome arms (Table S1). The deletion breakpoints were verified via PCR and subsequent Sanger sequencing (Fig S1A). An F1 male carrying a deletion of both *4930567H17Rik* coding sequences in cis *(4930567H17Rik*^ΔCDS/Y^) was bred to a C57BL/6J female to generate *4930567H17Rik*^ΔCDS/+^ female mice. *4930567H17Rik*^ΔCDS/+^ females were backcrossed to C57BL/6J males to generate *4930567H17Rik*^ΔCDS/Y^ mice, which were used for all experiments. *4930567H17Rik*^ΔCDS/Y^ mice used in the described experiments were backcrossed to C57BL/6J for >7 generations.

To generate mice lacking the *Mageb5* gene family, zygotes from *Mageb5*^ΔArm/+^ females crossed to *Mageb5*^ΔArm/Y^ mice [12] were injected with Cas9 protein (50 ng/μl), an oligonucleotide donor (10 ng/μl), and dual sgRNAs (30 ng/μl) targeting a ∼900 bp deletion of *Mageb5* (Table S1). These injections resulted in two independent *Mageb5*^ΔArmΔCDS/Y^ lines, ^“^L1” carrying a 860bp deletion and “L2” carrying a 400 bp deletion. The deletion breakpoints of the two lines were verified via PCR and Sanger sequencing (Fig S1B). F1 females with both the *Mageb5* palindrome arm and coding sequence deleted in cis were bred to C57BL/6J males to generate *Mageb5*^ΔArmΔCDS/+^ female mice. *Mageb5*^ΔArmΔCDS/+^ females were backcrossed to C57BL/6J males for >10 generations to generate *Mageb5*^ΔArmΔCDS/Y^ mice, which were used for all experiments.

Both *4930567H17Rik*^ΔCDS/Y^ and *Mageb5*^ΔArmΔCDS/Y^ mice transmitted the *4930567H17Rik* and *Mageb5* coding sequence deletions through the germline and no changes in overall health were observed due to off-target effects of CRISPR or as a consequence of the deletions. All mice used in these studies were between 3-7 months of age. *4930567H17Rik*^ΔCDS/Y^ and *Mageb5*^ΔArmΔCDS/Y^ mice were directly compared to wild type littermates (*4930567H17Rik*^+/Y^ and *Mageb5*^+/Y^ mice) in all experiments allowing for the minimization of the effects of genetic background and age. If wild-type littermates were not available, then age-matched controls were used. Because both *Mageb5*^ΔArmΔCDS/Y L1^ and *Mageb5*^ΔArmΔCDS/Y L2^ mice were able to be maintained easily (had normal breeding), *Mageb5*^ΔArmΔCDS/Y L1^ were used for experiments presented in this work unless otherwise specified. Cages were kept on ventilated racks at 72°F, 30-70% humidity, on a 12hr:12hr light: dark cycle in a specific-pathogen free room. Cages were monitored daily by husbandry personnel and changed every two weeks. Mice were given water and fed Lab Diet 5008 food *ad libitum*. Adult mice were sacrificed by CO_2_ asphyxiation followed by cervical dislocation and pups were sacrificed by decapitation in compliance with ULAM standard procedures in euthanasia. The Institutional Animal Care and Use Committee of the University of Michigan approved all animal procedures (PRO00009403) and all experiments followed the National Institutes of Health Guidelines of the Care and Use of Experimental Animals.

### Genotyping

Genotypes of *4930567H17Rik*^ΔCDS/Y^ and *Mageb5*^ΔArmΔCDS/Y^ mice were determined via PCR on DNA samples collected from 1-2mm tail snips. Tails were digested in 50mM NaOH for 20 minutes at 95°C and briefly vortexed to dissolve tissues. 50µl of Tris HCl (pH 6.8) was added to neutralize NaOH and samples were centrifuged at 13,000 rpm for 30 seconds [16]. PCR was performed with *Taq* DNA polymerase (New England Biolabs) per manufactures instructions. To verify genotypes of *4930567H17Rik*^ΔCDS/Y^ and *Mageb5*^ΔArmΔCDS/Y^ mice, we used primers flanking the coding sequence of each gene (primers 1-5 Table S2). For the *Mageb5* lines, we used primers flanking the *Mageb5* palindrome arm to verify the deletion of the palindrome arm, as previously described [12] (primers 6,7 Table S2).

### Reverse Transcriptase-PCR

Total testis RNA was extracted using Trizol (Life Technologies) according to the manufacturer’s instructions. ∼10µg of total RNA was DNase treated using Turbo DNase (Life Technologies) and reverse transcribed using Superscript II (Invitrogen) using oligo (dT) primers to generate first-strand cDNA. RT-PCR was performed on adult testis cDNA preparations with primers residing in the single exon coding sequence of *4930567H17Rik* (primers 3,8 Table S2), and with intron-spanning primers for *Mageb5* (primers 9,10 Table S2). Primers to the round spermatid-specific gene *Trim42* (primers 11,12 Table S2) served as a positive control [8]. To control for genomic DNA contamination, a reaction lacking reverse transcriptase was performed in parallel.

### RNA-Sequencing

Testis RNA was extracted from three *4930567H17Rik*^ΔCDS/Y^ and three *Mageb5*^ΔArmΔCDS/Y L1^ mice, along with three wild-type littermate controls from each line, and DNase treated as described above. Total RNA quality was assessed using the Tapestation 4200 (Agilent) (minimum DV200 value of greater than 30% and a minimum concentration of 3.32ng/µl). RNA used in this study had RIN values ranging from 6.1-8.9. Ribo-minus (RNaseH-mediated) stranded RNA-seq libraries with indexed adaptors were generated (New England BioLabs). Final libraries were quantitated by Kapa qPCR using Kapa’s library quantification kit for Illumina sequencing platforms (Kapa Biosystems, catalog # KK4835). Pooled libraries were subjected to 150 bp paired-end sequencing according to the manufacturer’s protocol (Illumina NovaSeq6000) giving an average of 50 million reads per sample. Bcl2fastq2 Conversion Software (Illumina) was used to generate de-multiplexed Fastq files. RNA-seq reads were pseudoaligned to the NCBI RefSeq gene annotation for the *Mus musculus* C57BL/6J (mm10) reference genome by Kallisto [17], using the default settings. Transcript per million (TPM) numbers were generated by Kallisto. The estimated number of RNA-seq reads aligning to each gene, as provided by Kallisto, were used as input to DESeq [18] to determine differentially expressed genes between *4930567H17Rik*^ΔCDS/Y^ and *Mageb5*^ΔArmΔCDS/Y^ and wild-type mice. All Illumina sequences can be found on NCBI’s sequence read archive under BioProject number: PRJNA748373 and accession numbers of SRR15198217 – SRR15198228.

### Testis Histology and Staining

Testes were fixed overnight in Bouin’s solution (Ricca Chemical, Arlington, TX). Following fixation, testes were washed in 5-10ml of 70% ethanol on a rotating tube holder at 4°C for 6-48 hours with three or more changes of ethanol to remove excess Bouin’s. Fixed testes were paraffin embedded, sectioned to 5μm, and stained with Periodic-Acid Schiff (PAS) and Hematoxylin. Testis sections were imaged on an Olympus BX61 equipped with an Olympus DP73 color camera. Specific germ cell populations were identified by their location within a tubule, nuclear size, and the nuclear staining pattern of chromatin [19].

### Testis to body weight ratio

To calculate testis/body weight ratio, total testis mass was divided by the total body mass taken at the time of euthanasia.

### Sperm counts and swim-up assay

Following dissection from the body cavity, the two cauda epididymis were dissected and nicked three times to allow sperm to swim out. Cauda were placed in 1ml of Human Tubal Fluid (HTF) (Millipore) at 37°C, and rotated in a 37°C incubator for 10 minutes. Cauda were removed and a 100μl aliquot was used for pre-swim-up baseline sperm counts. The remaining portion of sperm in HTF was then removed and placed in a new tube using wide bore tips. Sperm were centrifuged for 5 minutes at 400x*g* and the supernatant discarded. The pellet was re-suspended in 1ml of fresh 37°C HTF and centrifuged for 5 minutes at 400x*g*. The supernatant was removed and 1ml of fresh 37°C HTF was carefully overlaid on top of the pellet. The tube was then placed at a 45° angle in a 37°C incubator for 1 hour; after which the top 800μl containing motile sperm was removed and placed in a new tube. All aliquots of sperm used for counting were diluted 1:10 in H_2_O and counted using a hemocytometer. Counts were performed blind with four technical replicates per mouse. Sperm counts were calculated by taking the average number of sperm from each of the four technical replicates per mouse. Percent motility was calculated by dividing the post swim-up count by the pre-count and multiplying by 100. All analyses between groups were performed with an unpaired two-tailed student’s t-test, unless otherwise noted.

### Fecundity and Fertility

Three *4930567H17Rik*^ΔCDS/Y^ and three *Mageb5*^ΔArmΔCDS/Y^ mice, and equal numbers of wild-type litter mate controls, were each repeatedly mated with two CD1 females. Litter size was recorded, and the sex of each offspring was determined with sex-genotyping PCR primers (primers13,14 Table S2) specific to the X- and Y-linked gene *Ube1* (Ubiquitin-like modifier activating enzyme, 1 as previously described [8].

### Sperm head morphology assessment

To assess sperm head morphology, 25μl of the pre-swim-up sperm aliquot above was placed on a slide and allowed to dry. Cells were fixed for 10 minutes in 500μl of 4% PFA (diluted in PBS). Slides were rinsed twice in PBS for 5 minutes and left to dry. Slides were stained with Vectashield with DAPI (Vector Laboratories) under a 22 × 40 mm cover slip and imaged using an Olympus UPlanSApo 100x oil objective on an Olympus BX61 equipped with a Hamamatsu Orca-ER camera and Excelitas X-Cite 120LED fluorescence illuminator. Nucleus detection and morphology assessment was performed using the default settings of the custom plugin “Nuclear_Morphology_Analysis_1.20.0_standalone” to the image analysis software ImageJ [20]. ∼100 sperm heads from each genotype were blindly selected, imaged, and input into the software. Default edge detection settings were used, and sperm heads were manually inspected to ensure all sperm were accurately detected and only sperm were selected by the software. Sperm from *Mageb5*^ΔArmΔCDS/Y^ mice were not originally oriented correctly so the top and bottom vertical border tag was placed manually for the dataset [21].

## Results

### Mouse *4930567H17Rik* is a highly diverged ortholog of human *HSFX*, while mouse *Mageb5* is a conserved ortholog of human *MAGEB5*

X-linked gene families associated with large palindromes can have orthologs in other species or be independently acquired [4]. To assess possible orthologs of mouse *4930567H17Rik* and *Mageb5* in humans, we compared their protein sequence via BLASTP and found mouse *4930567H17Rik* is orthologous to human Heat Shock Transcription Factor X linked Member 3 (*HSFX3*) and mouse *MAGEB5* is orthologous to human MAGEB5 (Fig 1A). We further examined whether the genomic regions between mouse and human are syntenic (i.e. if they share flanking orthologous genes). In mouse, *4930567H17Rik* is flanked by the genes *Iduronate 2-sulfatase* (*Ids*) and *Transmembrane Protein 185a* (*Tmem185a*) which also flank *HSFX3* in humans. Interleukin 1 receptor accessory protein-like 1 *(Il1RAPl1) and Aristaless related homeobox* (*Arx*) flank *Mageb5* in both human and mouse (Fig 1A). This data supports that *4930567H17Rik* and HSFX3 and *Mageb5* and MAGEB5 are orthologous between human and mice.

**Figure 1.**
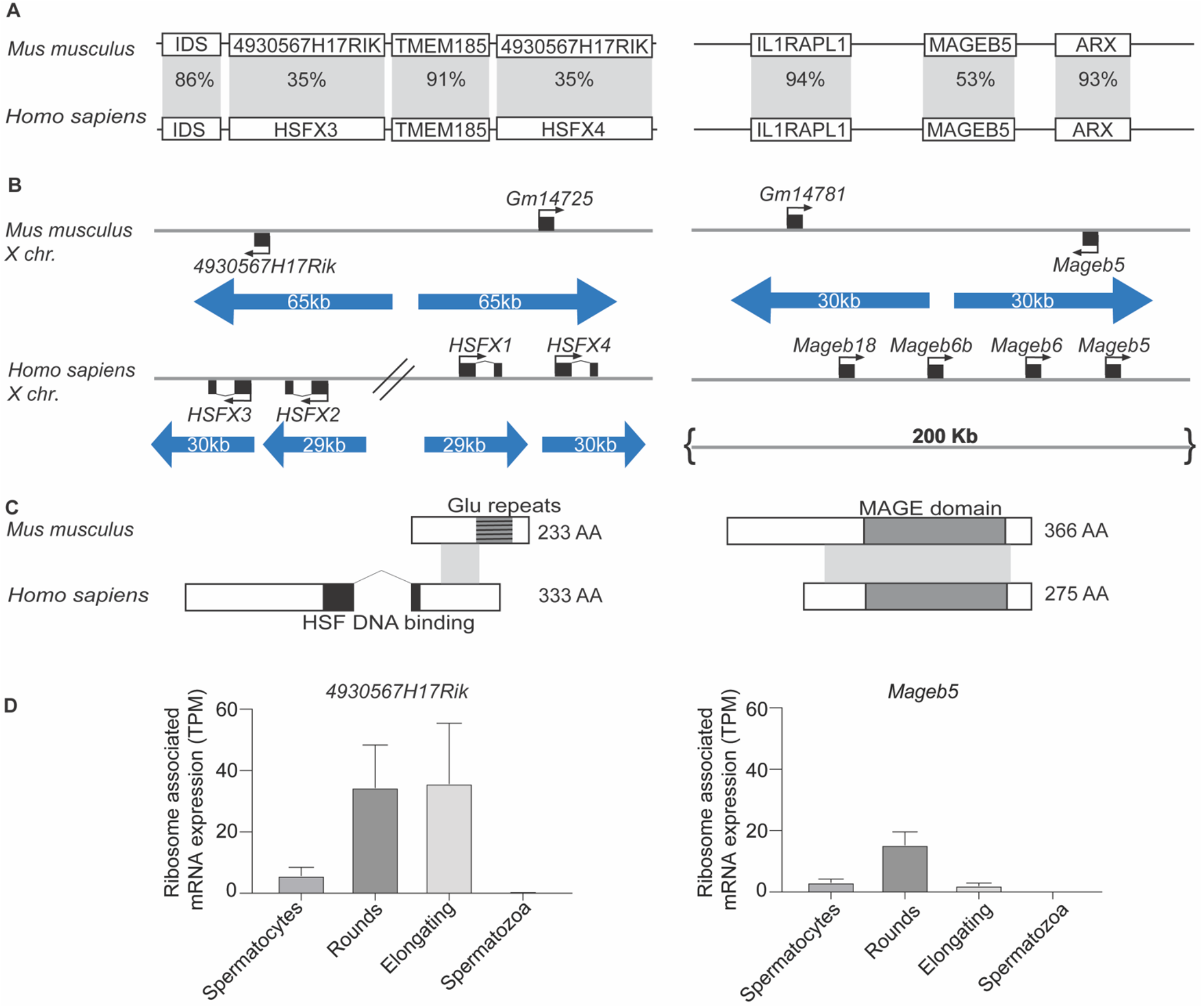
*4930567H17RIK* and *MAGEB5* share orthologs between mouse and humans. (A) Syntenic region of *4930567H17RIK* and *MAGEB5* between mouse and human with percent amino acid identity in the shaded region. *4930567H17Rik* shares 35% amino acid identity (across 18% of the protein) with *HSFX3. Mageb5* shares 53% amino acid identity (across 81% of the protein) with *MAGEB5* (B) Palindrome structure of the regions containing *4930567H17Rik, HSFX, Mageb5*, and *MAGEB5*. Palindrome arms are represented as blue arrows. *HSFX* is amplified compared to *4930567H17Rik. Mageb5* does not share palindrome structure between mouse and human. (C) Intron-exon structure for *4930567H17Rik*/*HSFX*, and *MAGEB5*/MAGEB5 showing protein domains (HSF DNA binding domain (black), glutamic acid repeats (dark grey), MAGE domain (light grey)) and amino acid sequence similarity (grey shading between species). (D) Reanalyzed ribosome profiling data for *4930567H17Rik* and *Mageb5* (data set taken from Wang, et. al. [24]). Ribosome association is most strongly seen in post meiotic cells for both *4930567H17Rik* and *Mageb5*.

In mouse, two gene copies of *4930567H17Rik* and *Mageb5* exist within palindromic sequences (Fig 1B), however, the copy number of both genes is different on the human X chromosome. In humans, *HSFX3* is present in four copies (annotated as *HSFX1-4)* and *MAGEB5* is inverted in a unique non-palindromic sequence. Human *MAGEB5* has additional neighboring *MAGEB* gene family copies, but none of the gene family members are within palindromes (Fig. 1B). BLASTP alignments and synteny of *4930567H17Rik/HSFX3* and *Mageb5/MAGEB5* suggest both X-linked gene families were present in the common ancestor of mouse and humans ∼80 million years ago (MYA), but diverged at the sequence level, as in the case of *4930567H17Rik*, or at the level of palindrome structure, as in the case of *Mageb5*.

### Mouse *4930567H17Rik* is a rapidly diverging protein-coding gene

To further understand how *4930567H17Rik* and *HSFX3* diverged, we compared the evolutionary dynamics and intron-exon structures of *4930567H17Rik* in rodents and *HSFX3* in primates. Previous studies have shown that both copies of *4930567H17Rik* (gene accession #’s NM_001033807, NM_001081476.1) exhibit rapid sequence divergence in rodents; having a K_a_/K_s_ value of 1.81 when compared across three *Mus* lineages with *R. norvegicus* as an outgroup [22]. We find primate *HSFX3* has a K_a_/K_s_ value of 1.20, indicating *4930567H17Rik* and *HSFX3* sequence is rapidly diverging in both rodents and primates. The rapid sequence divergence of *4930567H17Rik* in rodents may have been facilitated by the loss of an exon. Other mammalian *HSFX3* orthologs, including human *HSFX3*, encode two exons, while *4930567H17Rik* encodes a single exon (Fig1C). HSFX3 has a DNA binding domain spanning the splice junction between exon 1 and 2. *4930567H17Rik* produces a predicted protein that shares amino acid identity only with HSFX3. While 4930567H17RIK lacks the DNA-binding domain, it does possess an expanded glutamic acid repeat at the C-terminus end (Fig 1C). Interestingly, it appears the first exon of *4930567H17Rik* in mice has been pseudogenized and remnants of the first exon can be found outside of the palindrome via sequence comparisons (Fig S2). RNA-seq data supports that the ancestral first exon of HSFX is not transcribed in mice and thus the second exon is the only remaining functional exon in *Mus musculus* (Fig S2).

The predicted annotation of *4930567H17Rik* is a long non-coding RNA [23]. However, we find evidence to support that *4930567H17Rik* is a protein-coding gene despite the loss of an exon and rapid sequence divergence. First, *4930567H17Rik* encodes a large open reading frame (233 amino acids) that is conserved across rodents (Fig 1C). Despite the rapid sequence divergence, it has not acquired new nonsense mutations typical of pseudogenes or long non-coding RNAs. Second, reanalysis of ribosome profiling data [24] from sorted cell types across spermatogenesis demonstrate that *4930567H17Rik* RNA is most highly associated with ribosomes in round spermatid and elongating spermatid cell populations indicating that *4930567H17Rik* mRNA is likely translated (Fig 1D). Overall, we conclude that the protein-coding sequence of *4930567H17Rik/HSFX3* is rapidly diverging across mammals, and the exon containing the HSF DNA-binding domain has been lost along the Mus lineage, after divergence with rat.

### Generation of mice lacking both copies of the palindrome associated gene families *4930567H17Rik* or *Mageb5*

To determine whether the *4930567H17Rik* and *Mageb5* gene families are necessary for male fertility, we generated complete null mutant mice for both *4930567H17Rik* and *Mageb5* by using CRISPR/Cas9 (Fig 2A). We deleted both *4930567H17Rik* and *Gm14725*, which are both protein-coding gene copies (mm10 chrX:70,394,006:70,394,659 and chrX:70,545,638:70,546,279), resulting in a null mutant mouse line (*4930567H17Rik*^ΔCDS/Y^), as evidenced by RT-PCR and RNA-seq (Fig 2C,D). For *Mageb5*, we utilized *Mageb5*^ΔArm/+^ mice, which already had one palindrome arm which contained *Mageb5* deleted [12]. We deleted the second copy of the *Mageb5*, annotated as *Gm14781*, by specifically targeting the remaining protein-coding gene copy of *Mageb5* in *Mageb5*^ΔArm/+^ mice (Fig 2A) to generate two null mutant *Mageb5*^ΔArmΔCDS/Y^ lines (L1 and L2). In L1 mice, ∼860bp were removed (mm10 chrX:91,634,446-91,635,305), while in L2 ∼430 bp were removed (mm10: chrX:91,634,446-91,634,878) (Fig. 2B, right), yielding PCR products of 166bp (L1) and 596bp (L2). The translational start site was removed in both lines. To assess if *Mageb5*^ΔArmΔCDS/Y^ L1 and L2 produced RNA, we performed RT-PCR, followed by Sanger sequencing. We find that *Mageb5*^ΔArmΔCDS/Y^ L1 yields a highly truncated RNA product (322bp) and L2 does not yield any detectable RNA product (Fig 2C). Sequencing of the 322bp cDNA from L1 reveals a small predicted peptide (< 80 amino acids) (Fig S3). RNA-seq analyses of L1 support a lack of mRNA expression for both copies of the *Mageb5* gene in *Mageb5*^ΔArmΔCDS/Y L1^ mice (Fig. 2D). Our results support the successful removal of ∼650bp of both copies of *4930567H17Rik* and ∼860bp and ∼430bp of *Mageb5*, and 30Kb of a palindromic arm containing *Mageb5*, to yield *4930567H17Rik*^ΔCDS/Y^ and *Mageb5*^ΔArmΔCDS/Y^ null mutant mice, respectively.

**Figure 2.**
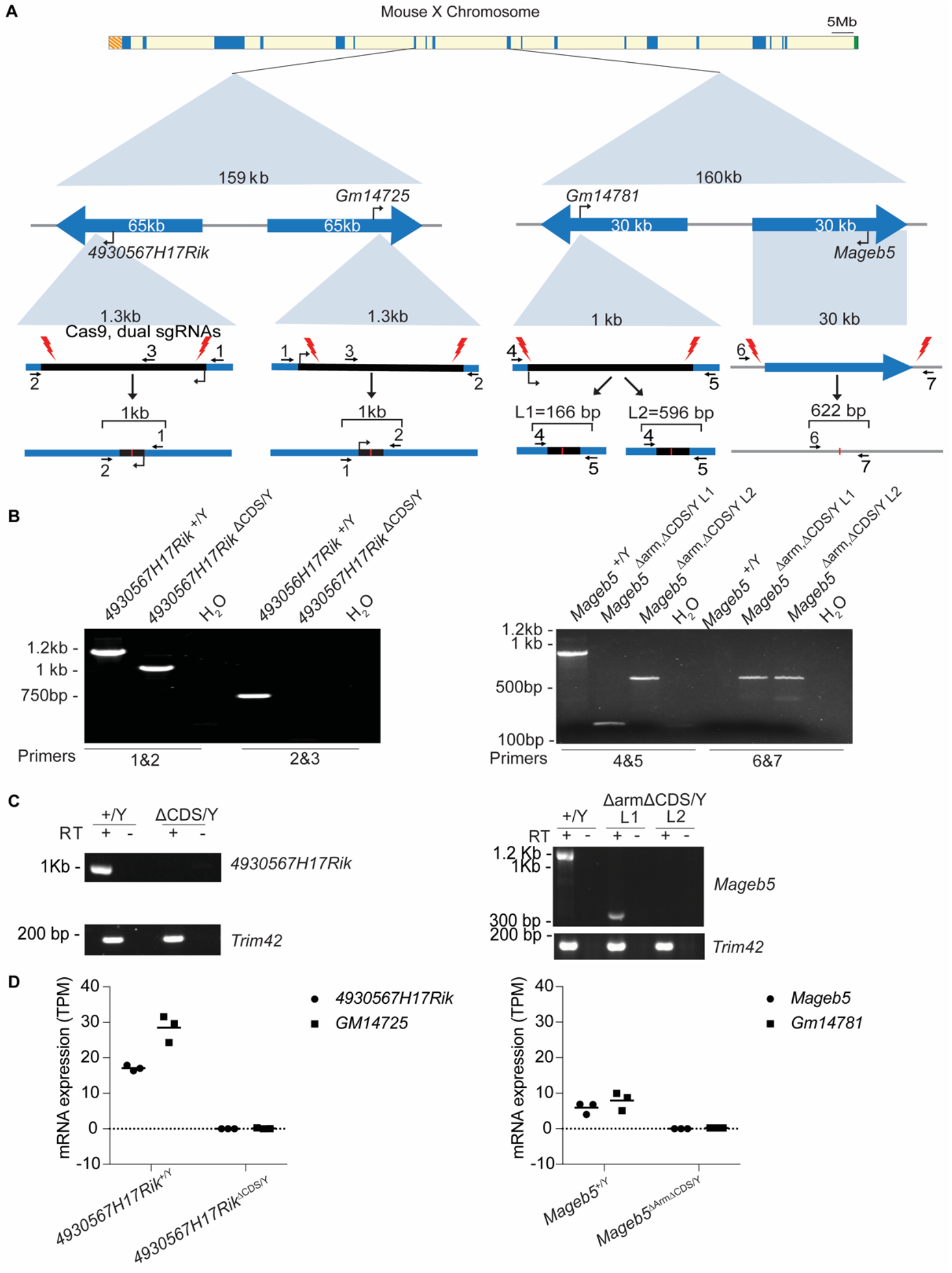
Creation of *4930567H17Rik*^ΔCDS/Y^, and *Mageb5*^ΔArmΔCDS/Y^ mice. (A) Top: Schematic of the mouse X chromosome showing singleton palindromes (blue) the pseudoautosomal region (green, right end), and centromere (orange, left end). Middle: Diagrams of the *4930567H17Rik* and *Mageb5* palindromes showing palindrome arms as blue arrows. Bottom: Coding sequence of *4930567H17Rik* and one copy of *Mageb5* with CRISPR cut sites (red lightning bolts) and genotyping primers (black arrows with numbers). (B) PCR genotyping of DNA from mutant and wild type *4930567H17Rik* and *Mageb5* mice. Numbered primers correspond to panel A. (C) RT-PCR of *4930567H17Rik* and *Mageb5* cDNA from one 4930567H17Rik line and *Mageb5* L1 and L2. L1 shows a product due to a small portion of the transcript still being produced. (D) RNA-seq data for *4930567H17Rik*^ΔCDS/Y^, and *Mageb5*^ΔArmΔCDS/Y L1^ mice. *GM14725* and *Gm14781* are the palindrome arm gene copies of *4930567H17Rik* and *Mageb5*, respectively.

### *4930567H17Rik* and *Mageb5* do not play major transcriptional regulatory roles during mouse spermatogenesis

We tested whether mouse 4930567H17RIK or MAGEB5 regulates transcription, since 4930567H17RIK is orthologous to a heat shock transcription factor, HSFX3, and MAGE proteins are known to regulate transcription [25, 26]. We therefore assessed whether mice lacking *4930567H17Rik* or *Mageb5* exhibit differences in transcriptional regulation. We performed whole testis RNA-seq to identify differentially expressed genes between *4930567H17Rik*^ΔCDS/Y^ or *Mageb5*^ΔArmΔCDS/Y^ mice and their wild-type littermate controls. We identified 21 and 27 differentially expressed (p-value <0.0001) genes for *4930567H17Rik*^ΔCDS/Y^ and *Mageb5*^ΔArmΔCDS/Y^ mice, respectively (Fig S4, Table S3). Consistent with our above analyses supporting a lack of *4930567H17Rik* and *Mageb5* expression, both copies of each gene family (*4930567H17Rik* and *Gm14725* and *Mageb5* and *Gm14781*) are the top two differentially expressed genes and are significantly downregulated in *4930567H17Rik*^ΔCDS/Y^ and *Mageb5*^ΔArmΔCDS/Y L1^ mice (Fig S4, Table S3). The limited number of differentially expressed genes, suggests these differentially regulated genes are involved in spermatogenesis suggests Thus, *4930567H17Rik* or *Mageb5* influence the transcription of a limited to a small collection of genes.

### *4930567H17Rik*^ΔCDS/Y^ and *Mageb5*^ΔArmΔCDS/Y^ mice display testis histology, sperm counts, and sperm motility similar to wild-type mice

To assess if *4930567H17Rik*^ΔCDS/Y^ and *Mageb5*^ΔArmΔCDS/Y^ mice exhibit defects in spermatogenesis or sperm biology, we examined if *4930567H17Rik*^ΔCDS/Y^ and *Mageb5*^ΔArmΔCDS/Y^ mice display defects in post-meiotic sperm development via testis histological sections. Periodicv acid-shift and hematoxylin (PAS+H) stained slides of testis tubules revealed that all stages of spermatogenesis were present and apparently similar to that of wild-type controls (Fig 3A) (Fig S4, Table S3), including a normal appearance of round spermatids, the cell type in which *4930567H17Rik* and *Mageb5* are predominantly expressed [3, 12]. In agreement with this finding, no significant differences in testis size (assayed via testis/body mass ratios) were detected (*4930567H17Rik*^ΔCDS/Y^= 0.006428±0.00086, *4930567H17Rik*^+/Y^=0.006167±0.00067, p=0.47; *Mageb5*^ΔArmΔCDS/Y^=0.007342±0.00050, *Mageb5*^+/Y^=0.007163±0.00069, p=0.54) (Fig 3B). Thus, in the absence of *4930567H17Rik* and *Mageb5* gene families, sperm development proceeds similar to that in wild-type testes.

**Figure 3.**
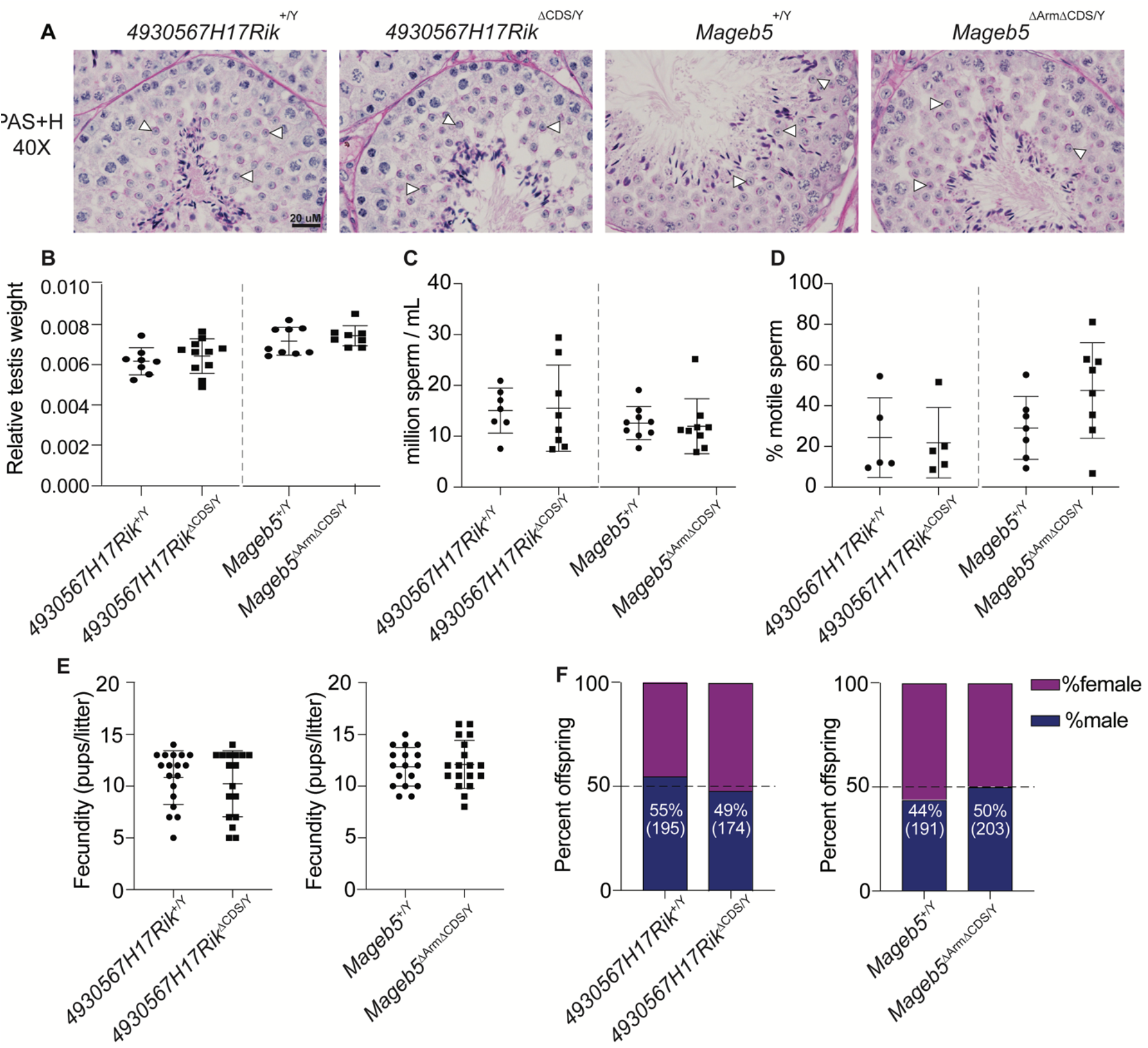
4*930567H17Rik*^ΔCDS/Y^ and *Mageb5*^ΔArmΔCDS/Y^ mice display testis histology, testis size, sperm counts, and motility similar to *4930567H17Rik*^+/Y^ and *Mageb5*^+/Y^ mice. (A) Periodic Acid-Schiff and Hemoxylin stained sections showing normal progression of spermatogenesis in *4930567H17Rik*^ΔCDS/Y^ and *Mageb5*^ΔArmΔCDS/Y^ mice. Arrows indicate presence of round spermatids. (B) Total testis weight normalized to total body weight (g). *4930567H17Rik*^+/Y^ n=8, *4930567H17Rik*^ΔCDS/Y^ n=11, *Mageb5*^+/Y^ and *Mageb5*^ΔArmΔCDS/Y^ n=9 (C) Sperm counts. Each point represents a single individual where 2 technical replicates were counted and averaged. *4930567H17Rik*^+/Y^ n=7, *4930567H17Rik*^ΔCDS/Y^ n=8, *Mageb5*^+/Y^ and *Mageb5*^ΔArmΔCDS/Y^ n=9.(D) Percent motile sperm. *4930567H17Rik*^+/Y^ and *4930567H17Rik*^ΔCDS/Y^ n=5, *Mageb5*^+/Y^ n=7 and *Mageb5*^ΔArmΔCDS/Y^ n=8. (E) Pups/litter from mating three males from each genotype with CD1 females. *4930567H17Rik*^+/Y^ n=18, *4930567H17Rik*^ΔCDS/Y^ and *Mageb5*^+/Y^ n=17 and *Mageb5*^ΔArmΔCDS/Y^ n=18 litters. (F) Sex genotyping performed on pups from litters shown in E. Number of male offspring are shown as percentage of total number of pups in parenthesis below. All comparisons in B-F were performed using an unpaired two-tailed student’s t-test between *4930567H17Rik*^+/Y^ vs *4930567H17Rik*^ΔCDS/Y^ and *Mageb5*^+/Y^ vs *Mageb5*^ΔArmΔCDS/Y^. All error bars represent mean with standard deviation.

We also assessed whether there were defects in sperm counts or sperm motility in *4930567H17Rik*^ΔCDS/Y^ and *Mageb5*^ΔArmΔCDS/Y^ mice. There were no differences in the number of sperm produced between *4930567H17Rik*^+/Y^ and *4930567H17Rik*^ΔCDS/Y^ and, *Mageb5*^+/Y^ and *Mageb5*^ΔArmΔCDS/Y^ mice (*4930567H17Rik*^ΔCDS/Y^=15.6±8.5×10^6^/ml, *4930567H17Rik*^+/Y^=15.0±4.4 ×10^6^/ml, p=0.88; *Mageb5*^ΔArmΔCDS/Y^ =11.9±5.4 ×10^6^/ml, *Mageb5*^+/Y^= 12.6± 3.3 ×10^6^/ml, p=0.78) (Fig 3C). Furthermore, sperm appeared to be equally motile regardless of genotype (*4930567H17Rik*^ΔCDS/Y^=22.91±17.99%, *4930567H17Rik*^+/Y^=24.95±20.37%, p=0.87; *Mageb5*^ΔArmΔCDS/Y^ 49.28±24.43%, *Mageb5*^+/Y^=30.14±16.09%, p=0.10) (Fig 3D).

### *4930567H17Rik*^ΔCDS/Y^ and *Mageb5*^ΔArmΔCDS/Y^ mice exhibit wild-type levels of fertility, fecundity, and sex ratio

To assess if *4930567H17Rik*^ΔCDS/Y^ and *Mageb5*^ΔArmΔCDS/Y^ mice sire fewer offspring, *4930567H17Rik*^ΔCDS/Y^ and *Mageb5*^ΔArmΔCDS/Y^ mice were mated to female CD1 mice. Both *4930567H17Rik*^ΔCDS/Y^ and *Mageb5*^ΔArmΔCDS/Y^ mice exhibited wild-type levels of fertility and fecundity, producing litters of equivalent size to wild-type controls (*4930567H17Rik*^ΔCDS/Y^=10.2±3.2 pups/litter, *4930567H17Rik*^+/Y^=10.8±2.6 pups/litter, p=0.55; *Mageb5*^ΔArmΔCDS/Y^=12.1±2.3 pups/litter, *Mageb5*^+/Y^=11.9±1.9 pups, p=0.77) (Fig 3E). Previous studies have found X-palindrome-associated genes can skew the offspring sex ratio [8], thus we genotyped the sex of all offspring. No sex ratio distortion was detected in the offspring of either *4930567H17Rik*^ΔCDS/Y^ or *Mageb5*^ΔArmΔCDS/Y^ mice (*4930567H17Rik*^ΔCDS/Y^ =49% male offspring, *4930567H17Rik*^+/Y^=55% male offspring, p=0.25; *Mageb5*^ΔArmΔCDS/Y^=50% male offspring, *Mageb5*^+/Y^= 44% male offspring, p>0.99, unpaired two-tailed t-test) (Fig 3F), suggesting mouse *4930567H17Rik* or *Mageb5* do not influence sex ratio distortion, autonomously.

### *4930567H17Rik*^ΔCDS/Y^ and *Mageb5*^ΔArmΔCDS/Y^ mice display altered sperm morphology

Defects in sperm morphology can still be present, despite wild-type levels of sperm production and motility, so we examined sperm morphology in *4930567H17Rik*^ΔCDS/Y^ and *Mageb5*^ΔArmΔCDS/Y^ mice. We analyzed multiple attributes of sperm morphology, including area, size of hook, and overall width of the sperm heads, using a custom plugin in ImageJ software [20]. We find that sperm from *Mageb5*^ΔArmΔCDS/Y^ mice were slightly more elongated than *Mageb5*^+/Y^ sperm (0.24 vs 0.16 p=<0.0001). Sperm from *4930567H17Rik*^ΔCDS/Y^ mice are larger overall (4739 vs 4420 square pixels, p<0.0001) and have slightly longer hooks (54 vs 52 pixels, p<0.0001) than *4930567H17Rik*^+/Y^ sperm (Fig 4A). The only additional statistical trend observed in morphology was the aspect ratio, the inverse of ellipticity, which was statistically different in *4930567H17Rik*^ΔCDS/Y^ (p<0.02) and in *Mageb5*^ΔArmΔCDS/Y^ mice (p<0.001) (Fig S6). Despite these differences, 4930567*H17Rik*^ΔCDS/Y^ and *Mageb5*^ΔArmΔCDS/Y^ mice are still fertile under laboratory conditions, suggesting that these morphological differences play a minor role in overall fertility despite these small morphological changes.

**Figure 4.**
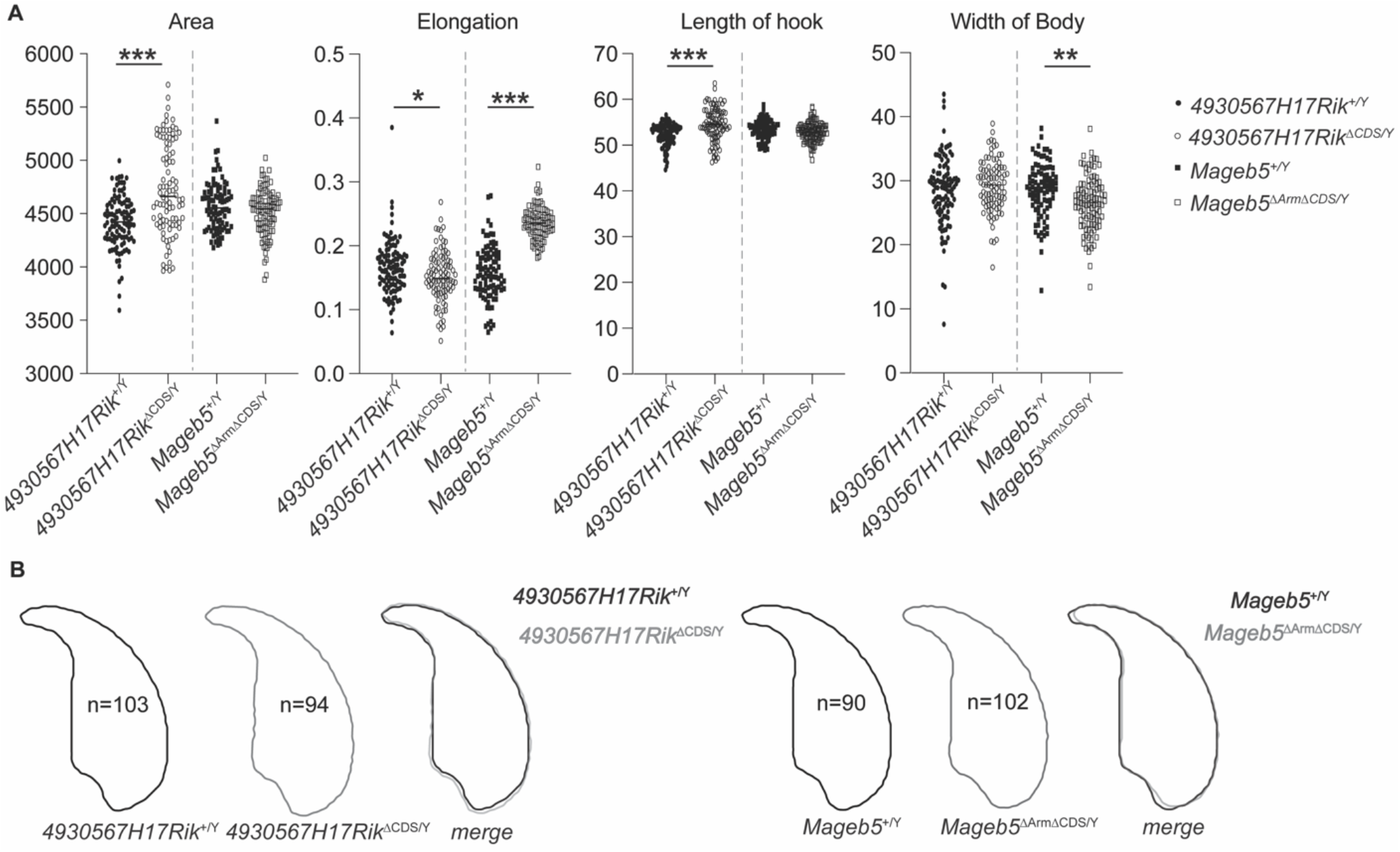
*4930567H17Rik*^ΔCDS/Y^ and *Mageb5*^ΔArmΔCDS/Y^ mice display altered sperm morphology. (A) Sperm morphology characteristics calculated from assessment of DAPI images processed with a custom plugin to ImageJ [20]. All data were compared using an unpaired two-tailed t-test between *4930567H17Rik*^+/Y^ vs *4930567H17Rik*^ΔCDS/Y^ and *Mageb5*^+/Y^ vs *Mageb5*^ΔArmΔCDS/Y^. * p<0.05 **p<0.001, ***p< 0.0001. Additional parameters are shown in supplementary figure S6. (B) Averages traces and shaded overlays of sperm head morphology from each genotype. Number of sperm assessed are shown inside each respective outline.

## Discussion

Our study addresses whether gene families harbored within large singleton X-palindromes are required for male fertility and spermatogenesis in mice. While null mutants of the *Slx* and *Slxl1* gene families harbored in X-palindrome arrays result in male infertility and defects in spermatogenesis [8], null mutants of the *4930567H17Rik* and *Mageb5* gene families in singleton X-palindromes do not. The absence of an overt reproductive phenotype in male mice lacking *4930567H17Rik* or *Mageb5* may in part be due to genetic redundancy. *4930567H17Rik* is related to heat shock transcription factors, which could have compensating family members. Indeed, *Hsf1* and *Hsf2* are expressed in post-meiotic cells [27, 28] and *Hsf2* is known to regulate post-meiotic X and Y palindromic genes [28] suggesting that *Hsf1* and *Hsf2* could compensate for the loss of *4930567H17Rik*. Similarly, *Mageb5* has eight X-linked [3] and one autosomal *Mageb* gene family members expressed in the testis that could potentially compensate for the loss of the *Mageb5* gene family. To better understand the spermatogenic role of *Mageb5*, and *Mageb* family members, removal of multiple *Mageb* family members may be necessary. For both *4930567H17Rik* and *Mageb5*, further studies investigating these possibly redundant genes could help elucidate the roles of *4930567H17Rik* and *Mageb5* in spermatogenesis. Furthermore, studies of *4930567H17Rik* orthologs in rats or primates, that still possesses HSF DNA-binding domains, could shed light on the ancestral function of *4930567H17Rik*.

Despite the lack of overt reproductive phenotypes in *4930567H17Rik*^*ΔCDS/Y*^ and *Mageb5*^*ΔArmΔCDS/Y*^ mice, differences in sperm head morphology suggest *4930567H17Rik* and *Mageb5* play a role in sperm development. Sperm head morphology analysis uses DAPI stained images of sperm to detect chromatin. *4930567H17Rik*^*+/Y*^ and *Mageb5*^*ΔArmΔCDS/Y*^ sperm had increased size and elongation, respectively, compared to wild type sperm (Fig 4). This finding may represent that sperm from these mice have a reduced level of chromatin compaction. Thus, *4930567H17Rik* and *Mageb5* may alter chromatin compaction during spermiogenesis and epididymal transit, a time in development when sperm chromatin compaction is dynamic [29]. Tracking the dynamics of sperm head morphology during development and epididymal transit may help further define the role of *4930567H17Rik* and *Mageb5* in sperm development.

Our current analyses of *HSFX* gene families and previous studies on *4930567H17Rik* [22] demonstrate that *HSFX* and *4930567H17Rik* sequences are rapidly diverging, suggesting the *4930567H17Rik/HSFX* gene family is under positive selection throughout mammals. The gene family’s presence within a X-linked palindrome may facilitate this rapid evolution in multiple ways. First, positive selection is known to be stronger for X-linked genes with male-beneficial functions, because of male sex chromosome hemizygosity [30]. Second, the second gene copy provide more substrate for new beneficial mutations upon which selection pressures can readily act [31]. Third, the second copy could relax constraint on palindromic genes and facilitate the acquisition of novel functions [31]. Fourth, any beneficial mutation arising in one gene copy could be readily spread to other gene copies in the palindrome through arm-to-arm gene conversion [22]. In the future, it will be important to connect how the rapid sequence divergence of *HSFX* and *4930567H17Rik* relates to their spermatogenic functions.

Large palindromic regions are challenging to study in mice and thus have not been a priority in large mouse knockout project consortiums [32-35]. CRISPR now enables the study of both X-palindromic structures and their associated gene families. Megabase-sized deletions of arrays of large palindromes demonstrate the necessity of large palindromes and their associated genes for male fertility [7-9, 12]. However, these studies cannot resolve whether the palindrome structure or the associated gene families are responsible for male infertility. Our study demonstrates how CRISPR can generate specific deletions of a single palindrome-associated gene family, while keeping the palindrome structure largely intact. Our study also improves our understanding of large X-palindrome-associated gene function, by demonstrating that individual X-palindrome associated gene families are dispensable for male fertility. Future studies using CRISPR to genetically dissect the importance of palindrome structures versus associated gene families in reproduction will provide a more complete understating of the importance of these large genomic regions and their implications in male fertility.

## Acknowledgements

We thank M. Arlt, C. Swanepoel, I. Mier, A. Lawson, D. de Rooij for editorial comments as well as D. de Rooij and M. Brogley for technical assistance. Additionally, we would like to thank the PATH-TMPRS, the Rogel Cancer Center Tissue and molecular Pathology Shared Resource (funding support NIH P30 CA04659229) for help with tissue sectioning and staining. We also would like to thank the transgenic animal model core at the University of Michigan for generation of our mutant mice, and the advanced genomics core at the University of Michigan for Sanger sequencing and RNA-seq.

## Notes

**Grant support:** This research was supported by the National Institutes of Health (R01HD094736 to JLM, and F32HD105375 to EAD), National Institute of Child Health and Human Development.

All authors declare that they have no conflicts of interest.

### Competing Interest Statement

The authors have declared no competing interest.

