## Supplemental Information for "X-linked palindromic gene families *4930567H17Rik* and *Mageb5* are dispensable for male mouse fertility"

Supplementary Information

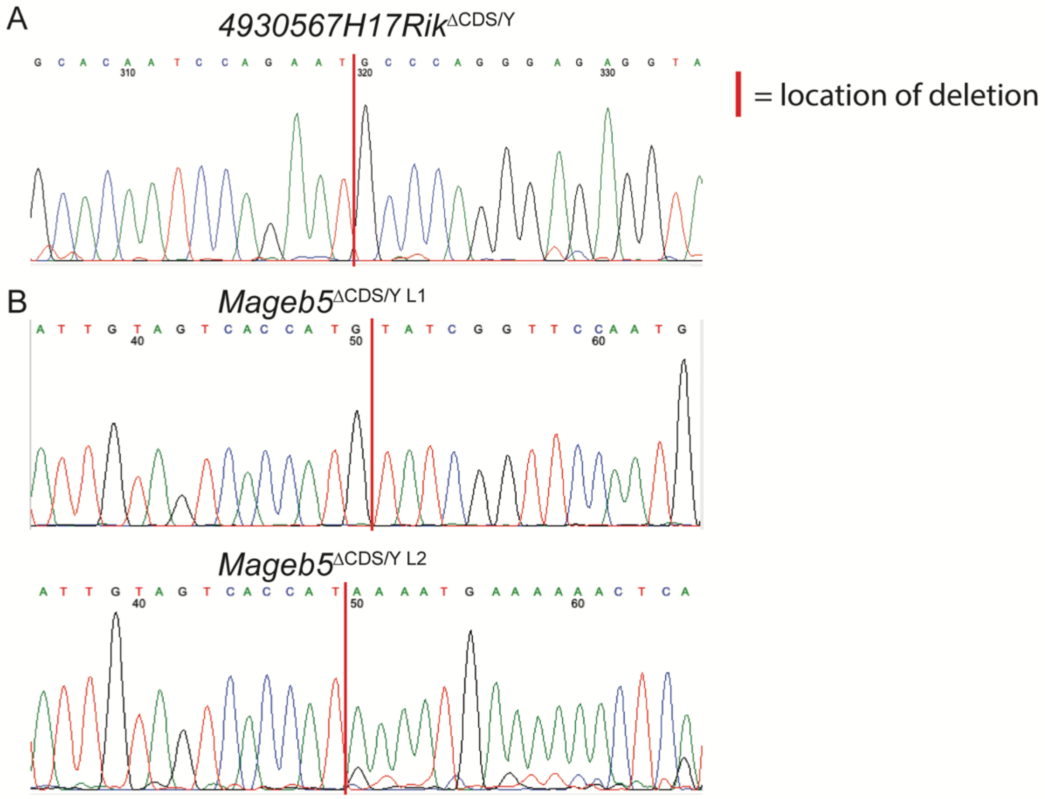


**Supplementary figure 1. Breakpoints in the coding sequences of *4930567H17Rik* and *Mageb5.*** Sanger sequencing results derived from PCR products showing breakpoints in *4930567H17Rik*$^{\Delta CDS/Y}$ *Mageb5*$^{\Delta Arm\Delta CDS/Y\ L1}$ and *Mageb5*$^{\Delta Arm\Delta CDS/Y\ L2}$ mice.

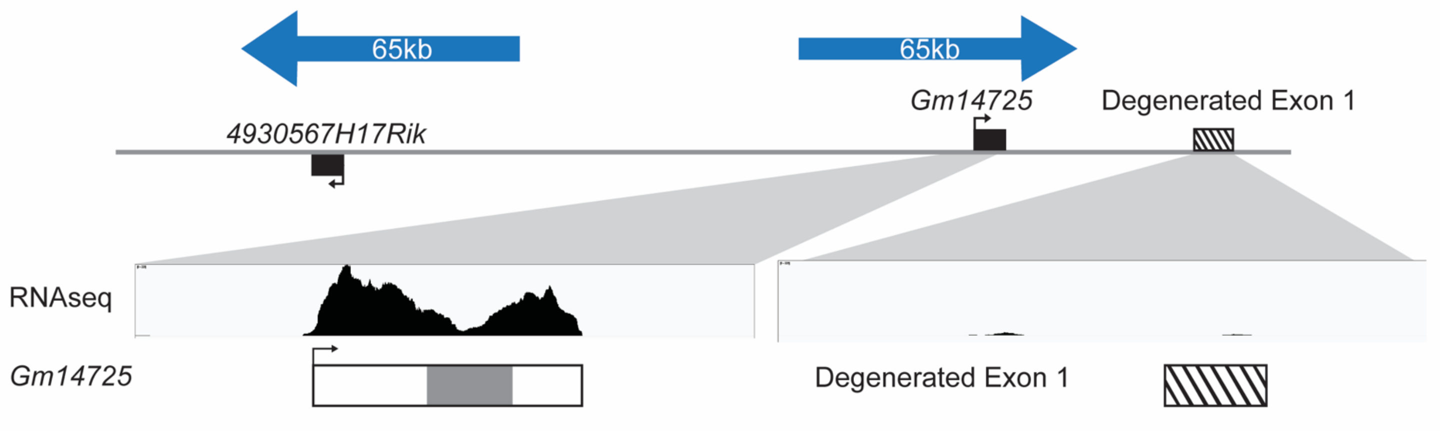


**Supplementary figure 2. Pseudogenization of a single exon of *4930567H17RIK* in mice.**
Top: RNA-seq data showing lack of transcription of the ancestral first exon (Degenerated Exon 1) of HSFX in mice. Bottom: Aligned representation of *Gm14725* (left) and the ancestral first exon (right), dark grey shading is a glutamic acid repeat region.

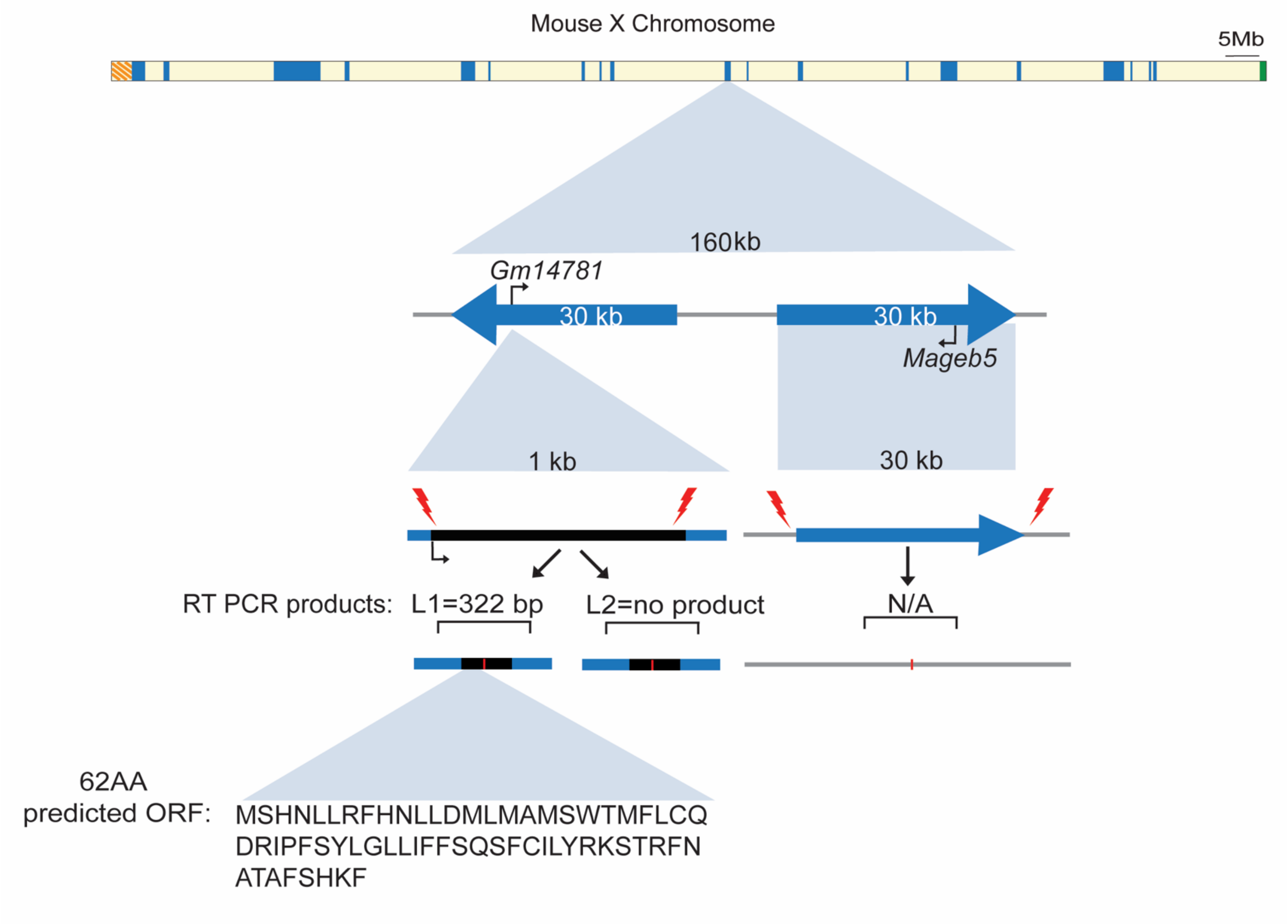


**Supplementary figure 3. RT-PCR analysis and predicted ORFs in *Mageb5*$^{\Delta arm,\ \Delta CDS/Y}$ mice.**

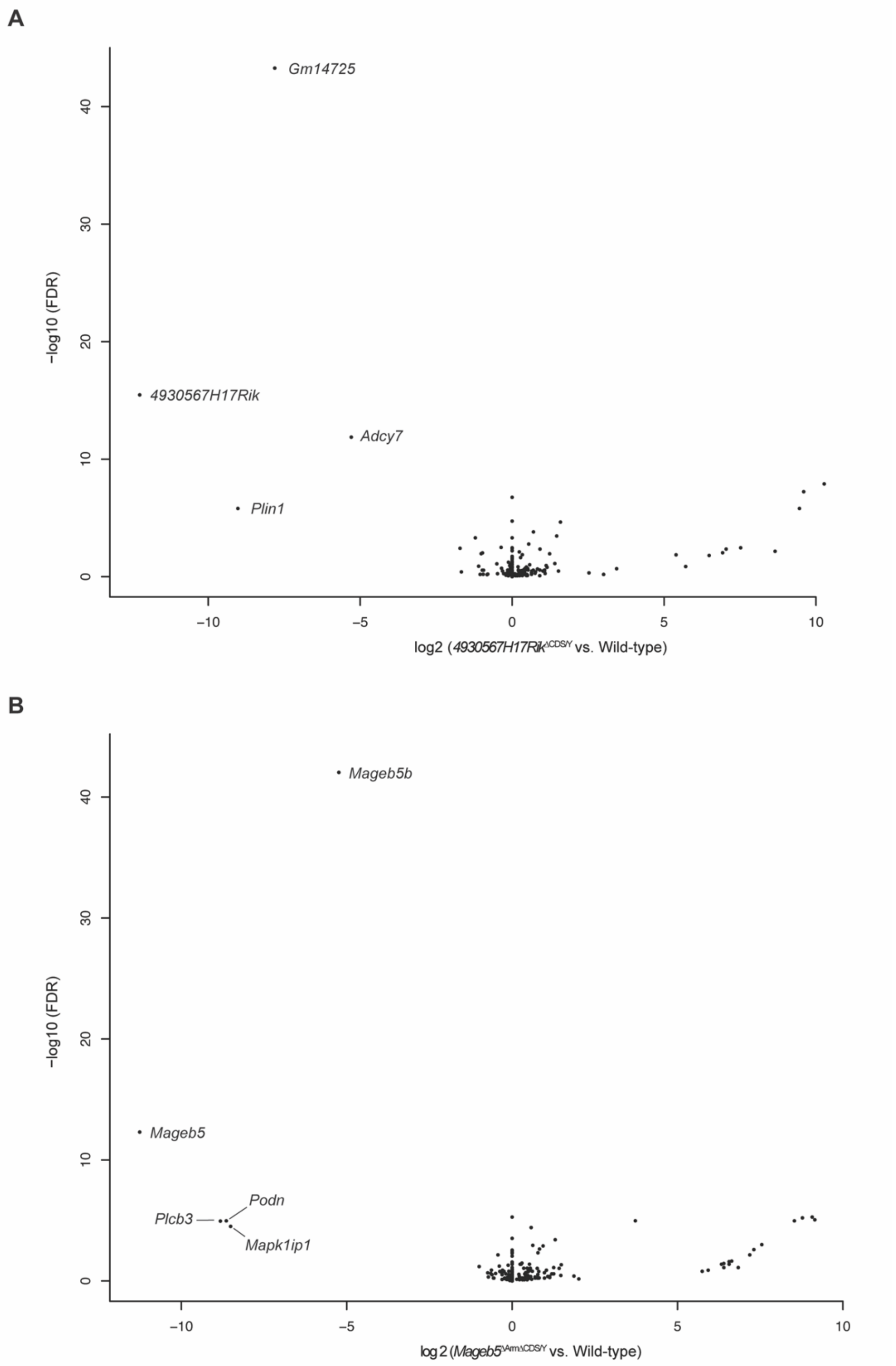


**Supplementary figure 4. Differential expression of genes in testes of *4930567H17Rik*$^{\Delta CDS/Y}$ and *Mageb5*$^{\Delta Arm \Delta CDS/Y}$ mice.** Volcano plots of differentially expressed genes (p-value <0.0001) from whole testis RNA-seq in *4930567H17Rik*$^{\Delta CDS/Y}$ (panel A) or *Mageb5*$^{\Delta Arm \Delta CDS/Y}$ (panel B) mice and their wild-type littermate controls.

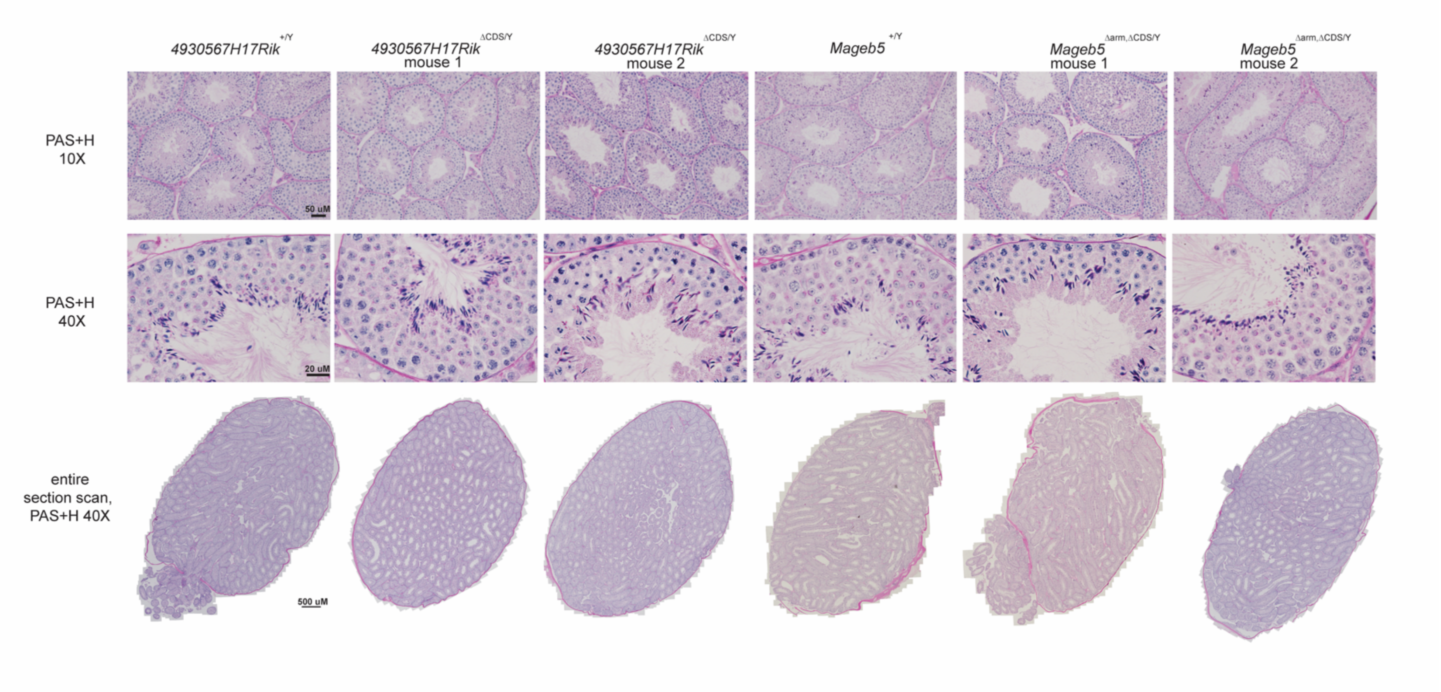


**Supplementary figure 5. Histological sections and whole section scans from *4930567H17Rik*$^{\Delta CDS/Y}$ and *Mageb5*$^{\Delta Arm \Delta CDS/Y\ L1}$ mice do not exhibit overt differences in spermatogenesis.** Sections were stained with periodic acid-Schiff and hematoxylin (PAS+H). Top row: 10X magnification, multiple tubules are displayed. Middle row: representative tubule from each individual used in the top row. Bottom row: scan of the entire testis section from each mouse.

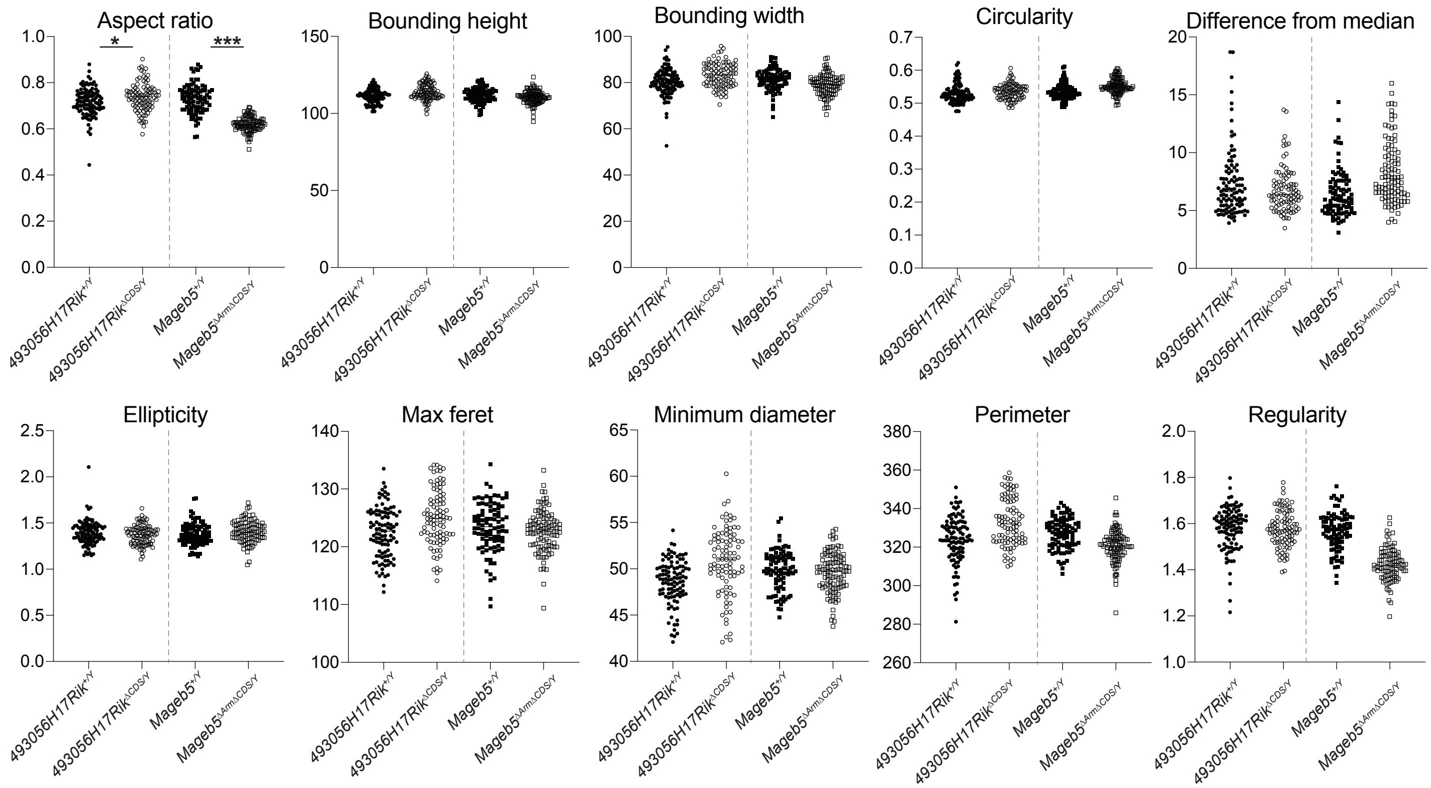


**Supplementary figure 6. Additional sperm morphology assessment parameters for *4930567H17Rik*$^{\Delta CDS/Y}$ and *Mageb5*$^{\Delta Arm\Delta CDS/Y}$ mice.** Characteristics calculated from assessment of DAPI images processed with a custom plugin to ImageJ from Skinner et al, 2019. * $p<0.05$ ** $p<0.001$ , *** $p< 0.0001$.

**Table S1.** sgRNA sequences used to generate *4930567H17Rik*$^{\Delta CDS/Y}$ and *Mageb5*$^{\Delta Arm\Delta CDS/Y}$ mice*

| Region | Sequence |
|---|---|
| 493Rik_gRNA_1 | CTGCACAATCCAGAATAGAC |
| 493Rik_gRNA_2 | TCCGTATGCT CTTCAGCCCA |
| 493Rik_CDSdel_ssODN | GAAAAAGGTAGTTATTCTCAGGCAGTACCTCTCCCTGGGCATTCTGGATTGTGCA GCATTAAACATAAATCAAGACATTC |
| Mageb5_gRNA_1 | TTGTAGTCACCATGCCC |
| Mageb5_gRNA_2 | CATTGGAACCGATAGCA |
| Mageb5_CDSdel_ssODN | TACTAGGTTCAACGCCACTGCCTTTAGCCATAAATTCTAGGACTTTCATCTTGGT AGTTTTAAGTAGAAGTA |

*Sequences used for arm deletions of the other gene copies of *4930567H17Rik* and *Mageb5* previously published in [1]

**Table S2.** Primers and sequences used to verify *4930567H17Rik*$^{\Delta CDS/Y}$ and *Mageb5*$^{\Delta Arm\Delta CDS/Y}$ mice and sex of pups for sex-ratio assays

| Figure : Primer number | Primer name | Sequence |
|---|---|---|
| 2A:1 | 493Rik_CDS_5'F | CCACCTCTTGAGGAATGGAA |
| 2A:2 | 493Rik_CDS_3'R | CAGGCAAGGAGGAGTGAGTC |
| 2A,C:3 | 4930567H17Rik_R | TCTGCATGGGTCGTATGA |
| 2A:4 | Mageb5_CDS F | TTGCCCTCTCATTATCTCCTACA |
| 2A:5 | Mageb5_CDS R | GCCACTGCCTTTAGCCATAA |
| 2A:6 | Mageb5_Arm F | GTTTGCAGAGTTGTGGACTGATAC |
| 2A:7 | Mageb5_Arm R | TGCATTGATCAAAAGGGAGA |
| 2C:8 | 4930567H17Rik_F | GGGCCTCTGAGACCACAT |
| 2C:9 | Mageb5 intron F | GGGAGAATCATCCACTTCTGA |
| 2C:10 | Mageb5 3' UTR | AAATCTCAAACGTGATGTTATAATTTC |
| 2C:11 | Trim42 F | GACTGTCTCAAGGCCTTC |
| 2C:12 | Trim42 R | CATGGCCATTGTGGAAAG |
| 13 | Ube1XY_F | TGGATGGTGTGGCCAATG |
| 14 | Ube1XY_R | CACCTGCACGTTGCCCTT |
